# SERCA function is impaired in skeletal and cardiac muscles from young DBA/2J mdx mice

**DOI:** 10.1101/2021.10.25.465805

**Authors:** Riley EG Cleverdon, Kennedy C Whitley, Daniel M Marko, Sophie I Hamstra, Jessica L Braun, Brian D Roy, Rebecca EK MacPherson, Val A Fajardo

## Abstract

The DBA/2J (D2) *mdx* mouse has emerged as a more severe model of Duchenne muscular dystrophy when compared to the traditional C57BL/10 (C57) *mdx* mouse. Here, we questioned whether sarco(endo)plasmic reticulum Ca^2+^-ATPase (SERCA) function would differ in muscles from young D2 and C57 *mdx* mice. In gastrocnemius muscles, both D2- and C57 *mdx* mice exhibited signs of impaired Ca^2+^ uptake, however, this was more severe in D2 *mdx* mice. Maximal SERCA activity was lowered only in D2 *mdx* gastrocnemius muscles and not C57 *mdx* muscles. Furthermore, in the left ventricle and diaphragm, Ca^2+^ uptake was impaired in C57 *mdx* muscles with lowered rates of Ca^2+^ uptake compared with C57 WT mice, whereas in muscles from D2 *mdx* mice, rates of Ca^2+^ uptake were unattainable due to the severe impairments in their ability to transport Ca^2+^. Overall, our study demonstrates that SERCA function is drastically impaired in young D2 *mdx* mice.

## INTRODUCTION

Duchenne muscular dystrophy (DMD) is an X-linked recessive muscle wasting disease, affecting approximately 1 in 5,000 boys worldwide (Mah et al., 2014; Mendell et al., 2012). Symptoms of DMD typically begin at three or four years of age and with no cure to date, patients usually succumb to cardiorespiratory complications from the disease by the third or fourth decade of life (Emery, 2002; Mavrogeni et al., 2015; Straub et al., 2016). Caused by the complete loss of dystrophin, a protein that normally connects the cytoskeleton and extracellular matrix to the sarcolemma, DMD is characterized by excessive membrane tearing and myofibre damage/degeneration (Blake et al., 2002). Loss of dystrophin is the initial insult of DMD, however there are several secondary consequences of the disease including chronic inflammation, oxidative stress, and calcium (Ca^2+^) overload (Tubridy et al., 2001). Combined, these processes ultimately lead to the pathological hallmarks of DMD which include degeneration, necrosis, and fatty and fibrotic replacement of muscular tissue (Shin et al., 2013).

Ca^2+^ is a potent signaling molecule largely responsible for excitation-contraction coupling (ECC) in skeletal and cardiac muscle. Two major Ca^2+^ regulatory proteins involved in muscle ECC are the ryanodine receptor (RyR) and sarco-endoplasmic reticulum ATPase (SERCA) pump, located in the sarcoplasmic reticulum (SR). RyR is responsible for the release of Ca^2+^ from the SR into the myoplasm, which initiates cross-bridge formation and force generation, whereas SERCA catalyzes the reuptake of Ca^2+^ into the SR to initiate muscle relaxation (Santulli et al., 2018; Tupling, 2004; Tupling, 2009). However, several studies have shown that if SR Ca^2+^ handling is dysregulated, Ca^2+^ can become chronically elevated in the myoplasm, leading to muscle damage and weakness. Indeed, impaired Ca^2+^ handling is a major part of the pathology in the muscles of patients living with DMD. The resulting Ca^2+^ overload exacerbates oxidative stress, inflammation, and cellular necrosis which in turn perpetuates dystrophic pathology (Robert et al., 2001; Turner et al., 1988). SERCA specifically has been implicated in dystrophic muscles, and several studies have shown that SERCA is impaired in skeletal and cardiac muscles from the murine C57BL/10ScSn (C57) *mdx* model of DMD (Gehrig et al., 2012; Schneider et al., 2013; Voit et al., 2017). This is in part due to elevations in reactive oxygen and nitrogen species (RONS) in *mdx* muscle that can damage the SERCA pumps (Gehrig *et al*., 2012). The SERCA pumps contain several residues that are susceptible to RONS mediated modifications such as nitrosylation and nitration, ultimately impairing their ability to transport Ca^2+^ (Viner et al., 1996; Viner et al., 1999a; Viner et al., 1997; Viner et al., 1999b).

The C57 *mdx* mouse, originally discovered in 1984 (Bulfield et al., 1984), has served as the dominant *in vivo* model to study cellular mechanisms and potential therapies for DMD. However, with a mild dystrophic pathology that is slow to progress, researchers have sought out alternative models that may better recapitulate human DMD (Swiderski and Lynch, 2021). For example, previous research has shown that SERCA pump function is more severely impaired in the *mdx*/utrophin double knockout (dko) mouse (Schneider *et al*., 2013; Voit *et al*., 2017) – a severe DMD mouse model where both dystrophin and its functional homolog utrophin are completely absent (Deconinck et al., 1997; Grady et al., 1997). This suggests that SERCA dysfunction progresses along with disease severity, whereby more severe models of DMD will display more severe impairments in SERCA function.

More recently, the traditional C57 *mdx* mouse was backcrossed onto a DBA/2J background, giving way to the *D2.B10-Dmd^mdx^*/J (D2 *mdx*) mouse (Coley et al., 2016). Unlike the C57 *mdx* mouse, the D2 *mdx* mouse presents with a juvenile onset and more severe DMD pathology, including pronounced muscle weakness and damage, degeneration, fibrosis, necrosis, and inflammation (Coley *et al*., 2016; Hammers et al., 2020; van Putten et al., 2019). The underlying mechanisms behind the greater pathology displayed in D2 *mdx* mice compared to C57 *mdx* mice are still under investigation, though a recent report has implicated elevated TGF-β signalling (Mázala et al., 2020). However, to our knowledge SERCA function has yet to be characterized in the D2 *mdx* mouse. Therefore, we examined SERCA function in 8-10-week-old D2 and C57 *mdx* mice. This cohort was selected based on previous studies showing that D2 *mdx* mice at this age display muscle weakness and pathology whereas the C57 *mdx* mice do not (Coley *et al*., 2016; Hammers *et al*., 2020; van Putten *et al*., 2019). We hypothesized that severity in pathology related to weakness observed in D2 *mdx* mice compared to the C57 *mdx* mice would be associated with greater impairments to SERCA function.

## RESULTS

### Muscle weakness and wasting in the C57 mdx versus D2 mdx mice

We first sought to verify the phenotypical weakness and atrophy previously reported in the D2 *mdx* mice. As expected, serum CK was elevated in both C57 and D2 *mdx* mice, denoted with a significant main effect of *mdx* genotype (Figure 1A). Moreover, gastrocnemius muscle and body mass were significantly lower in the D2 strain, indicating that mice on the D2 background are generally smaller. Interestingly, a significant interaction between background strain and *mdx* genotype showed that body mass was lower in D2 *mdx* vs D2 WT mice (Figure 1B and C). In contrast, there were no differences in body mass with C57 *mdx* and C57 WT mice. When normalized to body mass, a significant main effect of background strain was found suggesting that the gastrocnemius:body mass ratio was lower in mice on the D2 background compareed with those on a C57 background (Figure 1D). However, there were no differences in comparisons between WT and *mdx* mice for either C57 or D2 strains. Holding impulse, or hangwire time normalized to body mass, was significantly lower across both C57 *mdx* and D2 *mdx* mice compared with their respective WT groups as denoted by a significant main effect of *mdx* genotype (Figure 1E).

**Figure 1:**
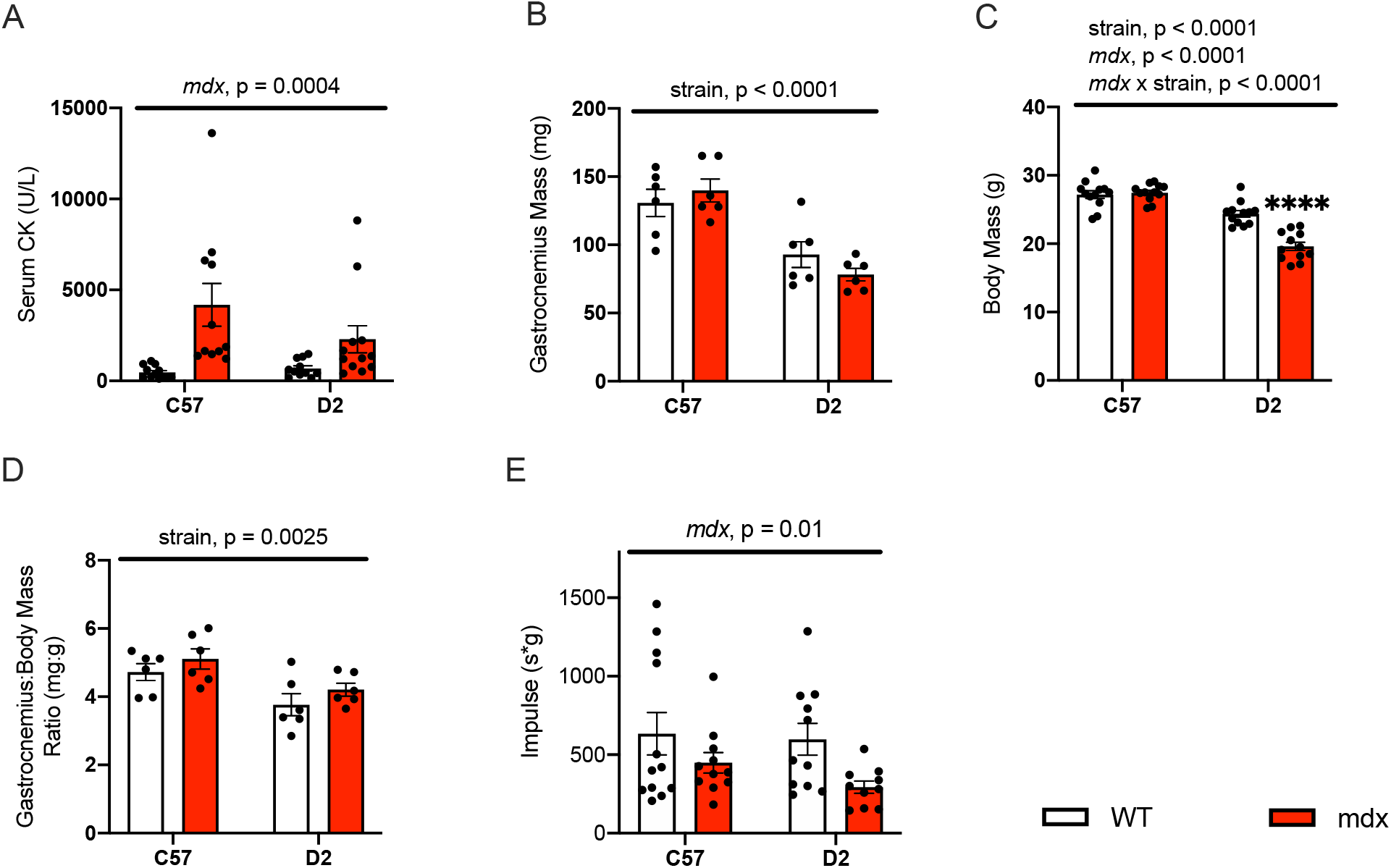
Creatine kinase, muscle/body mass and impulse: A) Serum creatine kinase activity expressed in units per litre. B) Gastrocnemius mass (mg), C) body mass (g), and D) gastrocnemius mass:body mass ratio (mg:g). E) Impulse expressed as hangwire time*body mass. A two-way ANOVA was used for all comparisons, n=12 per group (except for B and D, n = 6 per group). Values are represented as mean ± SEM. Main effects and interactions expressed in bars over the graph, \*\*\*\**p* < 0.0001 using a Sidak’s post-hoc test.

### Metabolic phenotyping of the C57 mdx and D2 mdx mice

Mice were housed in the Promethion metabolic cage system (48 hour periods) to examine cage activity, energy expenditure and various behavioural characteristics. Our data shows that D2 *mdx* mice were less ambulant in their cages compared with D2 WT mice with a main effect of *mdx* genotype measued across the light, dark and total daily phases (Figure 2A). However, we did not observe any differences in cage activity in C57 *mdx* mice compared with C57 WT mice, which is consistent with earlier-onset muscle weakness and wasting previously found in D2 *mdx* mice (Coley *et al*., 2016; Hammers *et al*., 2020; van Putten *et al*., 2019). Along with cage activity, we found that D2 *mdx* mice, but not C57 *mdx* mice had significantly elevated energy expenditure when compared to their respective WT groups (Figure 2 C and D). This was denoted as a significant main effect of *mdx* genotype in D2 mice (Figure 2C). Interestingly, this increase in daily energy expenditure in D2 *mdx* mice was met with a significant reduction in daily food intake compared with D2 WT mice (Figure 2E). We did not observe such changes in C57 *mdx* vs C57 WT mice (F). We also did not observe any changes in water intake across any experimental groups (Figure 2G and H). The Promethion metabolic cage system also allows for certain behavioural analyses including time spent in or touching its ‘home’ – a habitat enclosure on a slightly elevated platform. Our results show that D2 *mdx* mice spent less time in the home compared with D2 WT mice, however, this did not reach statistical significance (Figure 2I). There were no differences detected between C57 *mdx* and C57 WT mice (Figure 2J). We did find that D2 *mdx* mice spent significantly less time interacting or touching the habitat compared with D2 WT mice (Figure 2K), however, this was not observed in C57 *mdx* and C57 WT mice (Figure 2L). As the home is in an elevated platform, we believe that this serves as an additional indication of muscle weakness, whereby the *D2-mdx* mice were not strong enough to enter the elevated enclosure.

**Figure 2.**
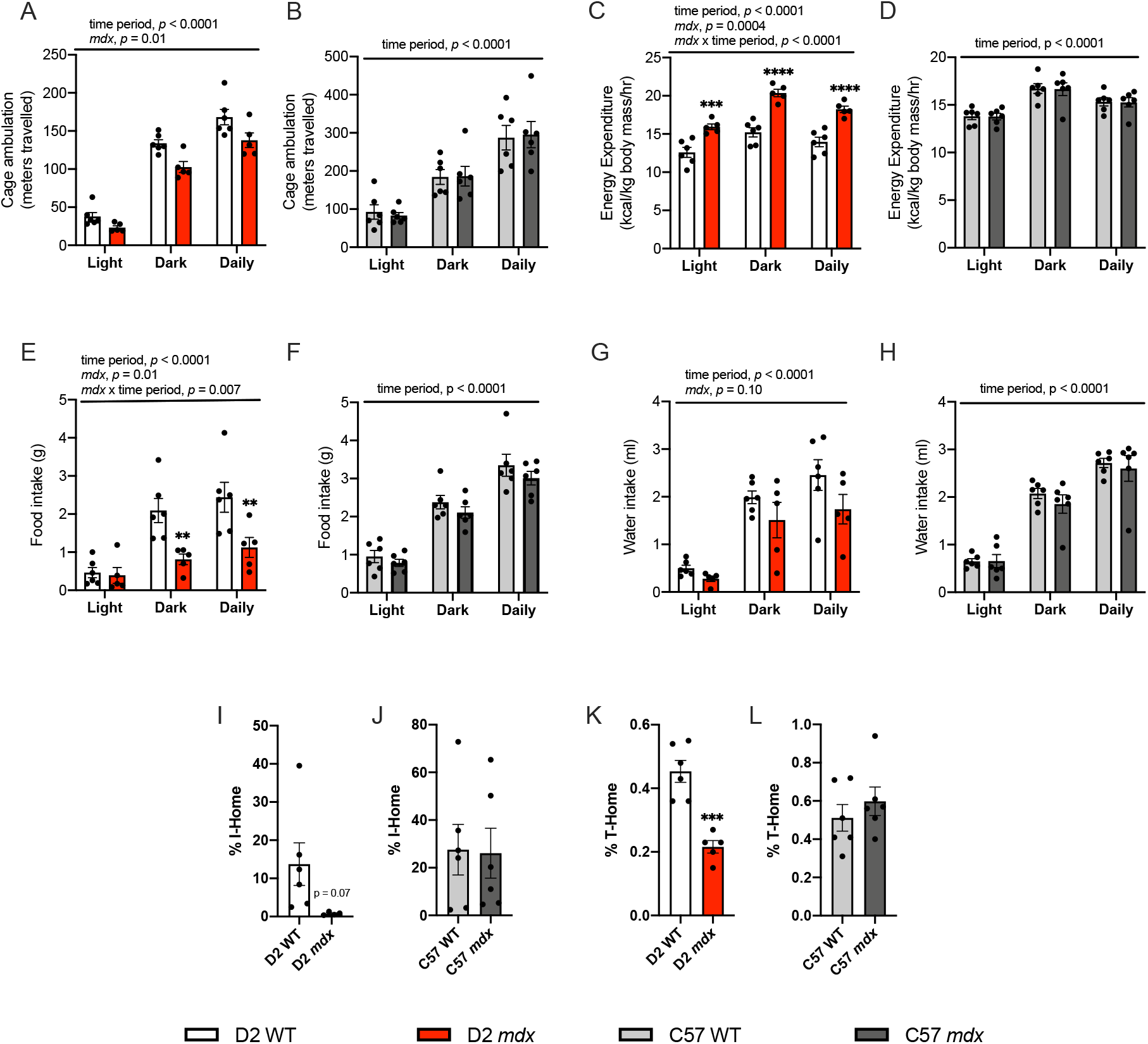
Cage activity, energy expenditure and cage behaviour: A) Daily cage ambulation in D2 mice and B) C57 mice expressed in metres travelled. C) Energy expenditure in D2 mice and D) C57 mice expressed in kcal/kg of body mass/hr. E) and G) Food (g) and water (mL) intake of D2 mice and F) and H) C57 mice. Percentage of time spent in (I-Home) and interacting with (T-Home) the mouse home for D2 mice I) and K) and C57 mice J) and L). All metabolic cage analyses were measured over a 48-hour period. Two-way repeated ANOVA examining the effect of time period, *mdx* genotype and their potential interaction (A-H). \*\**p* < 0.01, *** *p* < 0.001, and \*\*\*\**p* < 0.0001 using a Sidak’s post-hoc test, n = 5-6 per group. Student’s t-test examined WT versus mdx (I-L), n = 5-6 per group. Values are represented as mean ± SEM.

### Ca^2+^ uptake and SERCA activity is impaired in D2 mdx mice but not C57 mdx mice

We then examined SERCA function in gastrocnemius muscles from C57 and D2 -WT and *-mdx* mice (Figure 3). Ca^2+^ uptake experiments revealed obvious impairments in both C57 and D2 *mdx* mice (Figure 3A). Traditionally, rates of Ca^2+^ uptake would be examined using tangent analysis at a free Ca^2+^ concentration of 1000-2000 nM (Fajardo et al., 2015). However, for both C57 *mdx* and D2 *mdx* mice we could not utilize such approach as the Ca^2+^ uptake curve failed to cross 2000 and 1000 nM, which is indicative of impaired SERCA function. Alternatively, we examine the area-under-the-curve (AUC) and total Ca^2+^ taken in during the 400 s protocol (Figure 3B and C). For both, we found a significant main effect of *mdx* genotype with elevated AUC and lowered total Ca^2+^ uptake, suggesting that both C57 and D2 *mdx* mice have impaired SERCA function. However, we also note a significant interaction with background strain and *mdx* genotype, that shows that the elevation in AUC and reduction in total Ca^2+^ uptake is far more prominent in D2 *mdx* muscles (Figure 3B and C).

**Figure 3:**
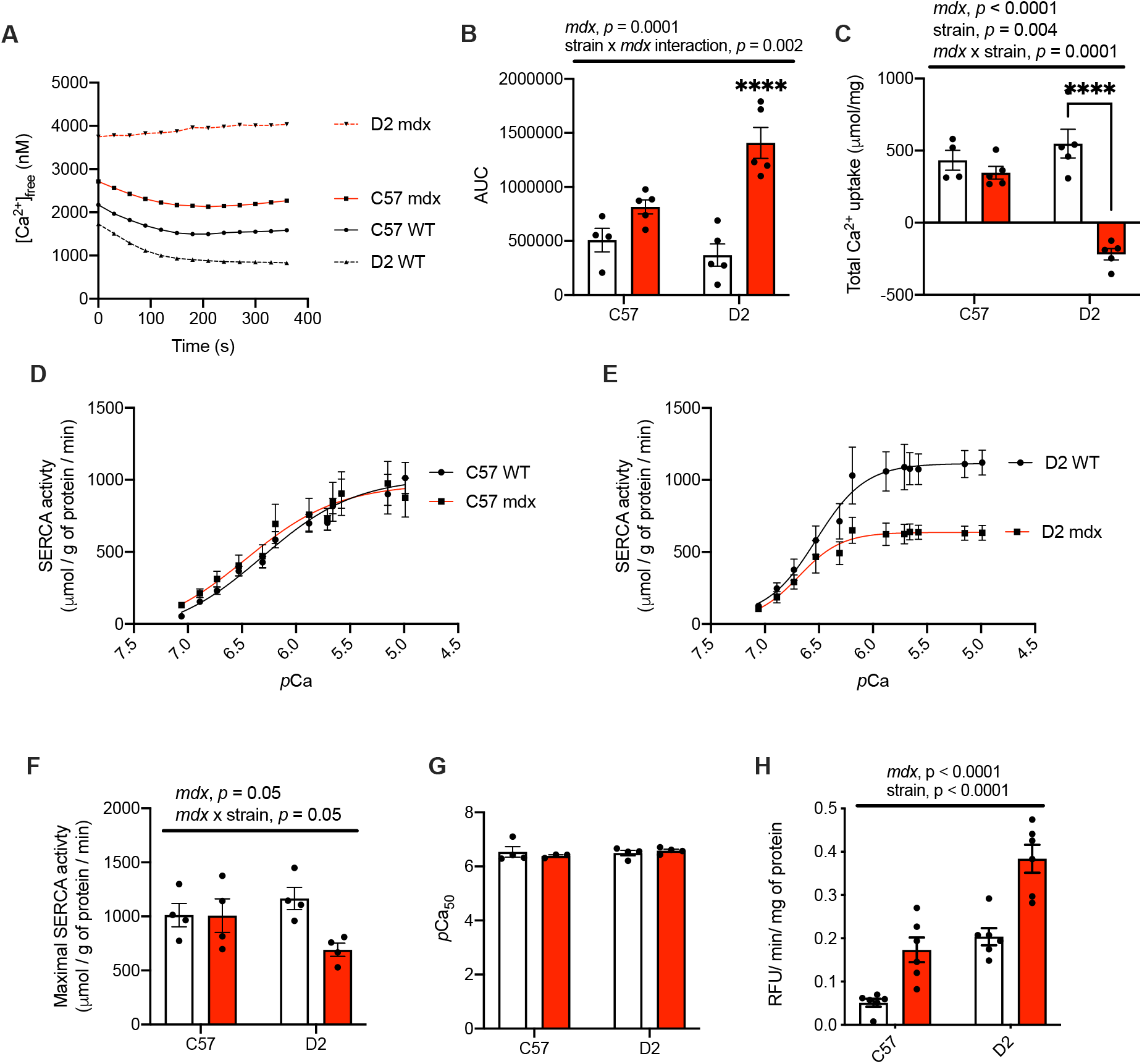
SR Ca^2+^ uptake/leak, SERCA and calpain activity in C57 and D2 -WT and *-mdx* gastrocnemius muscle: A) Ca^2+^ uptake curves from gastrocnemius homogenates. B) Area under the curve (AUC) of Ca^2+^ uptake curves. C) Total Ca^2+^ uptake normalized to wet tissue weight. D & E) Ca^2+^-dependent SERCA activity curves in D2 and C57 -WT and *-mdx* gastrocnemius muscles. F) Maximal SERCA activity expressed as μmol/g of protein/min and G) *p*Ca_50_ of SERCA activity, n = 4. H) Calpain activity in gastrocnemius muscles from D2 and C57 -WT and *-mdx* gastrocnemius muscles expressed as RFU min/mg of protein. Two-way ANOVAs were performed with a Sidak’s post-hoc test, \**p* < 0.05, \*\**p* < 0.01, \*\*\**p* < 0.001, and \*\*\*\**p* < 0.0001 n= 5-6. Main effects and interactions expressed in bars over the graph. Values are represented as mean ± SEM.

To further explore the impairments in SERCA function, we then measured Ca^2+^-dependent SERCA activity in the gastrocnemius muscle (Figure 3D and E). SERCA activity-*p*Ca curves show a selective impairment in D2 *mdx* gastrocnemius muscles; however, two-way ANOVA analysis did not reveal a significant interaction (p = 0.06, Figure 3F). No changes in SERCA’s apparent affinity for Ca^2+^ (*p*Ca_50_) were found (Figure 3G). Nonetheleless, our findings show that SERCA function is impaired in C57 and D2 *mdx* muscle, however, it the level of impairment is more prominent in the D2 *mdx* gastrocnemius muscle. To determine a potential pathological consequence of Ca^2+^ dysregulation calpain activity was measured in the gastrocnemius. We found that calpain activity was significantly higher in the C57 and D2 *mdx* muscles compared with C57 and D2 WT, as denoted by a significant main effect of *mdx* genotype (Figure 3H). In addition, a significant main effect of background strain indicates that D2 mice (WT and *mdx*) had greater calpain activity compared to C57 mice (Figure 3K). Furthermore, planned comparisons between D2 *mdx* and C57 *mdx* showed that calpain activity was significantly higher in D2 *mdx* mice *(p* = 0.0006, Student’s t-test).

### Elevated RONS may contribute to impaired SERCA function

To examine the potential mechanisms explaining the drastic imapirments in SERCA function found in D2 *mdx* muscles, we performed western blotting of SR Ca^2+^ handling proteins (Figure 4). Our results showed that SERCA1 content was upregulated in both C57 and D2 *mdx* mice with a significant main effect of *mdx* genotype (Figure 4A and B). A similar main effect was found with SERCA2 content, however, a significant interaction between *mdx* genotype and background strain indicated that the increase in SERCA2 was most prominent in D2 *mdx* muscle (Figure 4A and C). Sarcolipin (SLN) is a known inhibitor of SERCA that is upregulated in *mdx* mice (Fajardo et al., 2018; Schneider *et al*., 2013) and was ectopically expressed in both the D2 *mdx* and C57 *mdx* gastrocnemius muscle. That is, WT gastrocnemius muscles from WT C57 and D2 *mice* do not typically express SLN protein, and SLN could only be detected via western blot only in the *mdx* mice (Supplementary Figure 1). Interestingly, SLN content in D2 *mdx* gastrocnemius muscles was significantly lower compared with C57 *mdx* gastrocnemius muscle (Figure 4D). We then examined RyR1 content, and found that in both C57 and D2 *mdx* muscles, RyR1 was lower compared with C57 and D2 WT, with a significant main effect of *mdx* genotype (Figure 4E).RyR1 content in D2 mice (both WT and *mdx*) appeared to be lower than that found in C57 mice, but this was not statistically significant (Figure 4E). Calstabin, the channel stabilizer of RyR1 did not have any significant changes in protein content (Figure 4F). We also investigated protein nitrosylation and nitration in the gastrocnemius under reducing conditions to determine levels of irreversible RONS modification, since elevated RONS can impair SERCA function (Viner *et al*., 1996; Viner *et al*., 1999a; Viner *et al*., 1997; Viner *et al*., 1999b). Our results show that *mdx* mice (both C57 and D2) have elevated RONS levels compared with WT, with a main effect of *mdx* genotype. Furthermore, a significant interaction between *mdx* genotype and background strain shows that the elevations in total nitrocysteine and and nitrotyrosine were most prominent in D2 *mdx* mice (Figure 4G and H).

**Figure 4:**
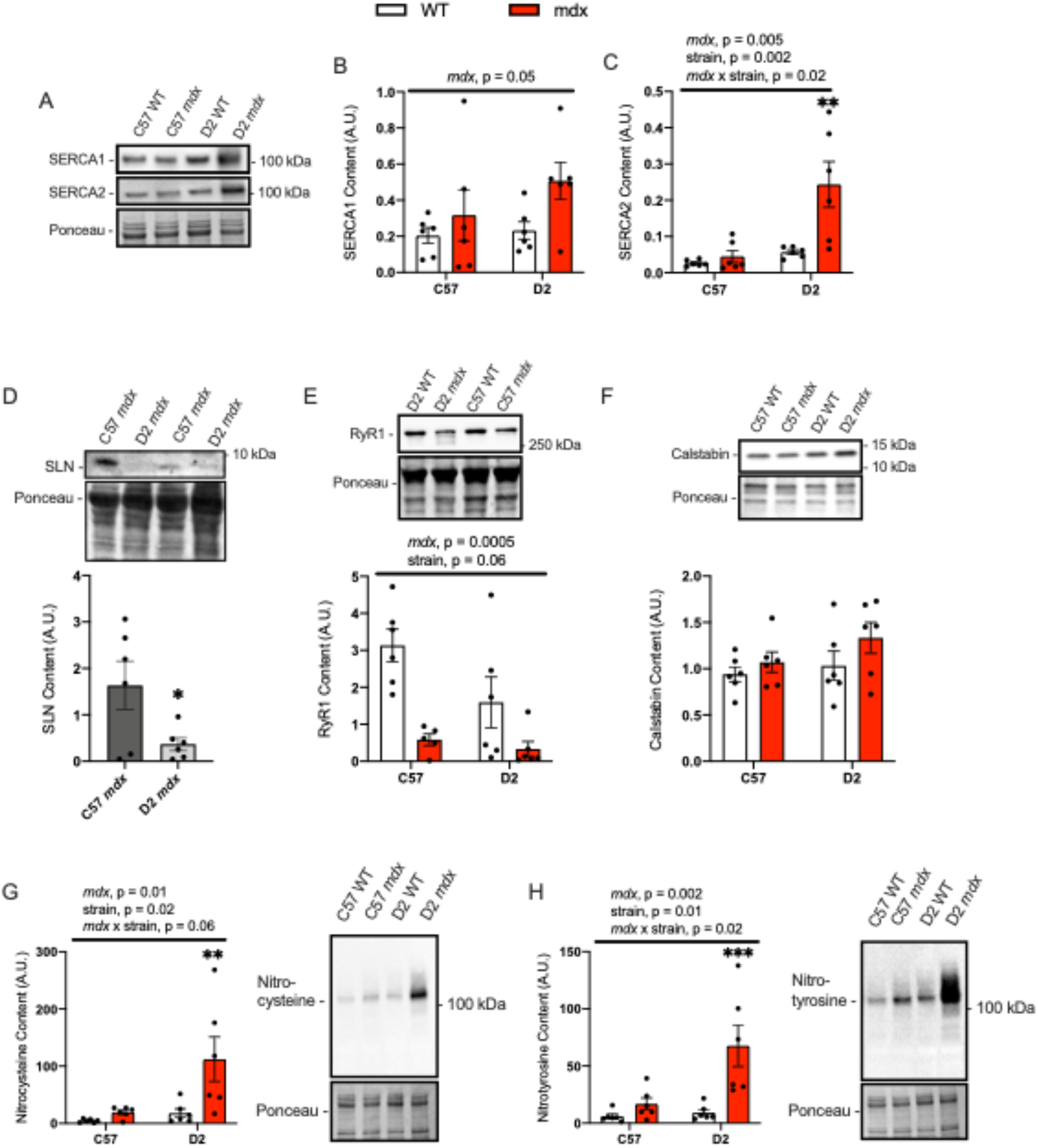
Western blot analyses of SR Ca^2+^ handling proteins total protein nitrosylation and tyrosine nitration: A) Representative Western blot images for SERCA1 and SERCA2 and their respective semi-quantificational analyses in gastrocnemius muscles from D2 and C57 -WT and *mdx* mice (B and C). D) SLN content in gastrocnemius muscles from C57 *mdx* and D2 *mdx* muscles. E) RyR1 and F) calstabin content in gastrocnemius muscles from D2 and C57 -WT and *-mdx*. G) Protein nitrosylation measured via western blotting for nitrocysteine-adducted protein. H) Protein tyrosine nitration measured via western blotting for nitrotyrosine-adducted protein. For B,C and E-H, a two-way ANOVAs were performed with a Sidak’s post-hoc test, \*\**p* < 0.01, \*\*\**p* < 0.001, \*\*\*\**p* < 0.0001 n = 6 per group. Main effects and interactions expressed in bars over the graph. For D a Student’s t-test was used, \**p* < 0.05, n = 6 per group. All protein contents are expressed as arbitrary units. Values are represented as mean ± SEM.

### Ca^2+^ uptake is also impaired in D2 mdx left ventricle and diaphragm muscles

Given the cardiorespiratory implications of DMD, we also assessed SERCA function in cardiac (left ventricle) and diaphragm muscles from C57 and D2 -WT and *-mdx* mice. Similar to our results with the gastrocnemius, we found that D2 *mdx* mice had severe decrements in Ca^2+^ uptake in both the cardiac and diaphragm muscles (Figure 5). In the cardiac muscle, Ca^2+^ uptake curves show a drastic impairment in Ca^2+^ transport in D2 *mdx* mice, which translated to a significant elevation in AUC when compared with D2 WT mice (Figure 5A and B). Though AUC was not different between C57 *mdx* and C57 WT mice, we were able to measure Ca^2+^ uptake rates at a free Ca^2+^ of 1000 nM. In doing so, we found a significant reduction in Ca^2+^ uptake rate in the LV from C57 *mdx* mice compared with C57 WT mice (Figure 5C). Similar findings were found in the diaphragm, where there were drastic impairments in Ca^2+^ uptake in the D2 *mdx* diaphragm that led to a significantly higher AUC compared with D2 WT mice (Figure 5D and F). Moreover, comparing Ca^2+^ uptake rates at 1000 nM, shows a significant difference between C57 WT and C57 *mdx* diaphragm muscles, with rates being lower in the latter (Figure 5F). Altogether, these results show that while Ca^2+^ uptake is indeed impaired in C57 *mdx* muscles, the impairment appears to be more prominent in D2 *mdx* muscles where rates of Ca^2+^ uptake were unattainable.

**Figure 5:**
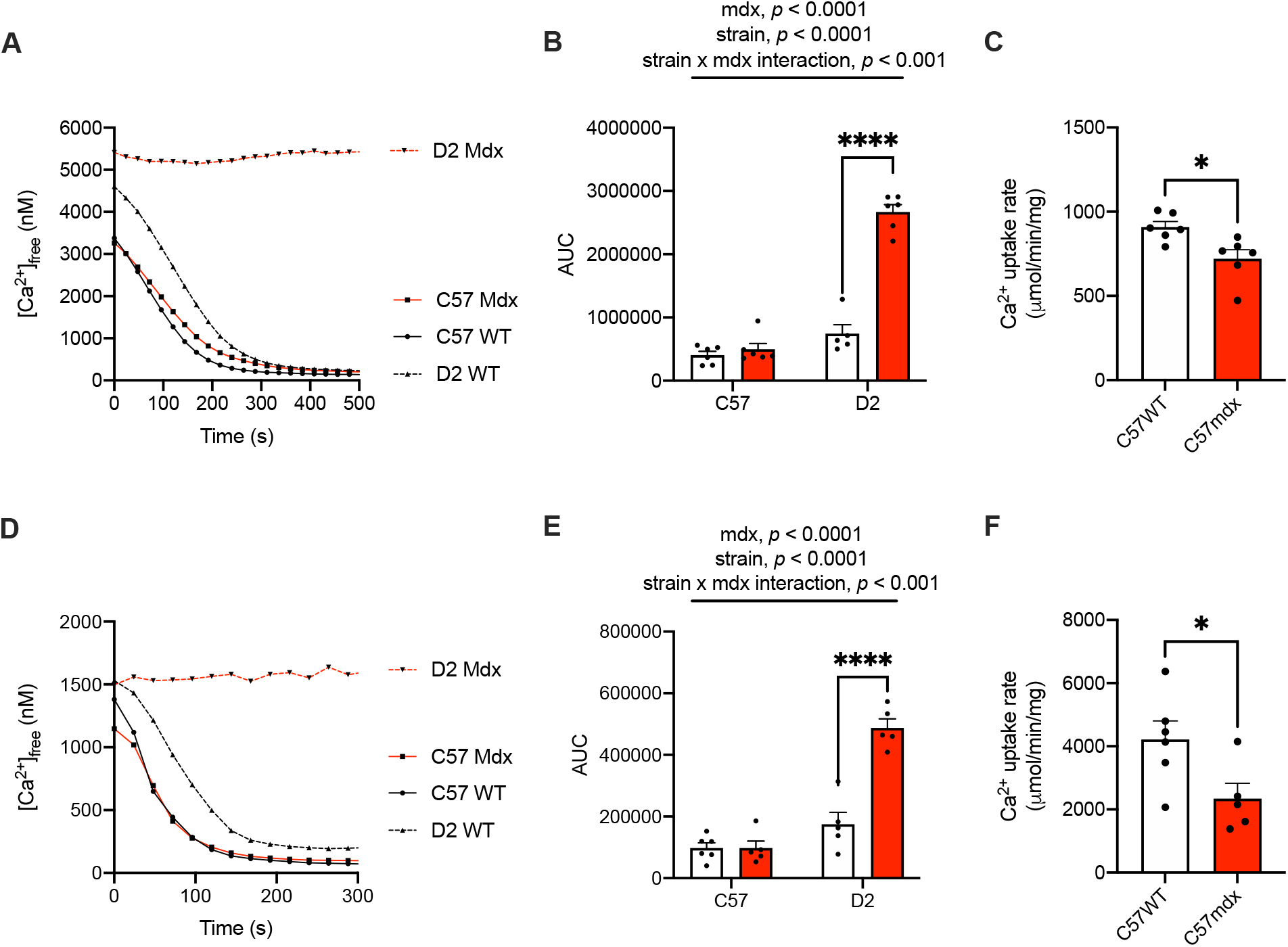
Ca^2+^ uptake in the cardiac and diaphragm muscles: A) Ca^2+^ uptake curves from left ventricle homogenates. B) AUC analysis in left ventricle homogenates. C) Ca^2+^ uptake rate in LV measured via tangent analysis at 1000 nM free [Ca^2+^]. D) Ca^2+^ uptake curves from diaphragm homogenates. E) AUC analysis in diaphragm homogenates. F) Ca^2+^ uptake rate in diaphragm muscles measured via tangent analysis at 1000 nM free [Ca^2+^]. For B and E, Two-way ANOVAs were performed with a Sidak’s post-hoc test, \*\*\*\**p* < 0.0001. For C and F, a Student’s t-test was used, \**p* < 0.05. Main effects and interactions expressed in bars over the graph. Values are represented as mean ± SEM, n = 5-6 per group.

## DISCUSSION

To our knowledge, this study is the first to compare SERCA function in both the D2 *mdx* mouse and C57 *mdx* mouse model. The D2 *mdx* mouse is known as a more severe DMD model with early onset weakness compared to the C57 *mdx* mouse. Impulse and measures of cage ambulation supported this showing that D2 *mdx* mice had early onset muscle weakness occurring at 8-10 weeks of age. Also, the D2 *mdx* mice had less interaction with the elevated mouse home enclosure compared to C57 *mdx* mice, suggesting that these mice were less able to jump onto this raised platform found within their cages. Interestingly, D2 *mdx* mice also had a hypermetabolic profile, where these mice had the highest levels of energy expenditure while ambulating the least, which we believe can contribute to the smaller body and muscle mass commonly observed in these mice. Perhaps inconsistent with our results, a previous study found that energy expenditure was elevated in 5 month old C57 *mdx* mice compared with WT mice (Strakova et al., 2018). We believe that this discrepancy could possibly be explained by age differences, as our C57 *mdx* mice were younger as compared to the mice used in the study by Strakova and colleagues.

To explain why the D2 *mdx* mice are much weaker at an earlier age, we explored the involvement of SERCA dysfunction. Our Ca^2+^ uptake results showed that D2 *mdx* muscles (gastrocnemius, cardiac and diaphragm) had obvious impairments in their ability to bring Ca^2+^ into the SR. This manifested as an elevated AUC during the uptake period. Importantly, several studies have shown that C57 *mdx* muscles exhibit impairments in SERCA function (Gehrig *et al*., 2012; Schneider *et al*., 2013; Voit *et al*., 2017). The findings presented in this study are consistent with this as C57 *mdx* gastrocnemius also had elevated AUC during the Ca^2+^ uptake protocol, and C57 *mdx* LV and diaphragm had significantly slower rates of Ca^2+^ uptake compared with C57 WT counterparts. However, the fact that Ca^2+^ uptake was so dramatically impaired in D2 *mdx* mice that rates of Ca^2+^ uptake at 1000-2000 nM free [Ca^2+^] could not be attained, in combination with lowered maximal SERCA activity, suggests that the impairment in SERCA function is most prominent in D2 *mdx* muscles. Altogether, our finding is consistent with a previous study showing that SERCA dysfunction proceeds on a continuum of disease severity, where more severe dystrophic models such as the dko mouse (Schneider *et al*., 2013) and the D2 *mdx* mouse have more severe impairments in SERCA function.

The alterations to SERCA pump function do not seem to be explained by differences in SERCA protein levels. Both SERCA1 and SERCA2 were upregulated in the gastrocnemius, which we believe is an adaptive response aimed at improving SR Ca^2+^ handling. Indeed, SERCA density is an important determinant of SR Ca^2+^ filling, at least in healthy conditions (Tupling, 2004). However, any compensatory increase in SERCA content/density was unsuccessful in the rescuing SERCA function. We next examined SLN expression. Although it is not typically expressed in murine gastrocnemius muscle, ectopic expression of SLN occurred in both *mdx* mouse models, with SLN protein being elevated to a greater extent in the C57 *mdx* mice compared to the D2 *mdx* mice. Studies have shown that SLN is upregulated along with disease progression with its expression being further upregulated in the dko mice compared with *mdx* mice (Schneider *et al*., 2013). Furthermore, knocking out SLN in mdx and dko mice improved dystrophic pathology suggesting that SLN may contribute to *mdx* pathology by improving SERCA function (Balakrishnan et al., 2022; Mareedu et al., 2021; Voit *et al*., 2017). However, there are discrepant results with SLN deletion studies also impairing *mdx* pathology (Fajardo *et al*., 2018) or not improving phenotype (data not shown in ref (Bal et al., 2021)). The reasons for these discrepant results are unclear, however, a point on hormesis was recently raised (Chambers et al., 2022). Since we found that SLN was less upregulated in the more severe D2 *mdx* mouse model with dramatic impairments in SERCA function, our results could suggest that SLN may not be a critical player in the SERCA function impairments and potentially *mdx* pathology in the D2 *mdx* mouse; however, this should be investigated further specifically in the D2 *mdx* mouse.

In addition, SLN is a known uncoupler of SERCA that uncouples Ca^2+^ transport from ATP hydrolysis (Bombardier et al., 2013; Smith et al., 2002). Thus, its upregulation in the D2 *mdx* mouse may contribute to the observed increase in daily energy expenditure. However, since we found that SLN levels were relatively higher in the C57 *mdx* vs D2 *mdx* mice, other potential mechanisms may be contributing to the increased daily energy expenditure found specifically in D2 *mdx* mice. One potential factor may be elevated oxidative/nitrosative stress as SERCA is susceptible to damaging RONS modifications such as nitrosylation and nitration (Viner *et al*., 1996; Viner *et al*., 1999a; Viner *et al*., 1997; Viner *et al*., 1999b). Consistent with our observations of a prominent elevation in total nitrocysteine and nitrotyrosine levels in D2 *mdx* muscles, it is plausible that RONS modifications could be contributing to the changes in SERCA function. That is, our results of elevated total protein nitrosylation and tyrosine nitration under reducing conditions are suggestive of irreversible RONS modifications that may be impairing the SERCA pump. Moreover, damage to the SR membrane itself from RONS may contribute to the drastic impairments in Ca^2+^ uptake found in D2 *mdx* muscles. While we note that maximal SERCA activity appeared to be lower in the D2 *mdx* mouse, there still was an appreciable amount of SERCA-mediated ATP hydrolysis. Combined with an obvious inability to bring Ca^2+^ into the SR, our results could suggest that the SERCA pumps have become highly inefficient in D2 *mdx* muscle – hydrolyzing ATP while bringing only minimal levels of Ca^2+^ into the SR. This uncoupling effect could potentially contribute to the elevated daily energy expenditure found in the D2 *mdx* mice.

We also determined the activation of calpain in D2 *mdx* gastrocnemius muscles. Calpain is a Ca^2+^-dependent proteolytic enzyme previously shown to be more active in *mdx* muscles (Gailly et al., 2007). To our knowledge, ours is the first study to demonstrate that rates of calpain activity are higher in the D2 background strain compared to C57, irrespective of the *mdx* genotype. Furthermore, planned comparisons between D2 *mdx* and C57 *mdx* mice specifically revealed a significant 2.2-fold increase in calpain activity in the former vs the latter (*p* = 0.0006 via Student’s t-test). This finding is important and adds mechanistic insight that may partly explain the worsened pathology in the D2 *mdx* mouse. It is interesting that the D2 WT mice already had elevated calpain activation compared with C57 WT. This apparent difference in background strain may provide additional insight as to why *mdx* mice on the D2 background are worse off than *mdx* mice on the C57 background. One explanation that is not yet well understood, is a potential crosstalk between calpain activation and the pro-fibrotic factor TGF-β. Previous studies have found that TGF-β1 upregulation in lung fibroblasts (Li et al., 2015) and atrial derived myocytes (Yeh et al., 2011) can increase calpain activation. This is interesting given recent findings showing that a polymorphism of LTBP4 in D2 WT mice greatly upregulates TGF-β (Heydemann et al., 2009; Mázala *et al*., 2020), suggesting that the increased TGF-β activity could increase calpain activity in the D2 WT mice. This combined with the impairments observed in SERCA function could contribute to elevated calpain in the D2 *mdx* mice.

In conclusion, our study clearly shows drastic impairments in SERCA function in skeletal and cardiac muscles from D2 *mdx* mice. Our results suggest that the early onset weakness and damage observed in young D2 *mdx* mice may be partly due to an inability to bring Ca^2+^ into the SR. Future studies should investigate whether improving SERCA function, potentially through cytotoxic protection against RONS may mitigate dystrophic pathology and improve physiological outcomes in the D2 *mdx* mouse.

## LIMITATIONS TO THE STUDY

Our study clearly shows that SERCA function is severely impaired early on in D2 *mdx* mice and may contribute to the severity of muscle weakness and damage. However, we acknowledge that our work is largely descriptive, and future studies aimed at improving SERCA function will determine its causal role, if any. Furthermore, the mechanisms leading to these impairments in SERCA function are still unknown, and though we associate them with elevated oxidative/nitrosative stress, SERCA-specific modifications should be assessed in the future. RyR Ca^2+^ leak is another major player in the Ca^2+^ disturbances found in C57 *mdx* muscle (Bellinger et al., 2009; Capogrosso et al., 2018). Our study is limited in that we did not examine RyR function. Finally, our study does not discern whether the elevated daily energy expenditure in D2 *mdx* mice is caused solely by increased muscle-based energy consumption. Other thermogenic organs such as brown and beige fat could be investigated in the future.

## ACKNOWLEDGEMENTS

This work was supported by an unrestricted Brock University internal grant and a Canada Research Chair Tier 2 in Tissue Plasticity and Remodelling to VAF. REGC and SIH were supported by a NSERC CGS-M, and JLB was supported by a CIHR CGS-M.

## AUTHOR CONTRIBUTIONS

**Riley EG Cleverdon**: Conceptualization, methodology, validation, formal analysis, investigation, writing – original draft, visualization

**Kennedy C Whitley**: Methodology, investigation, formal analysis, writing – review & editing

**Daniel M Marko**: Methodology, investigation, formal analysis, writing – review & editing

**Sophie I Hamstra**: Investigation, formal analysis, writing – review & editing

**Jessica L Braun**: Investigation, formal analysis, writing – review & editing

**Brian D Roy**: Resources, writing – review & editing

**Rebecca EK MacPherson**: Resources, supervision, writing – review & editing

**Val A Fajardo**: Conceptualization, supervision, project administration, funding acquisition, investigation, formal analysis, visualization, writing – original draft

## STAR Methods

### RESOURCE AVAILABILITY

#### Lead contact

Further information and requests for resources and reagents should be directed to and will be fulfilled by the lead contact, Val A. Fajardo.

#### Materials availability

This study did not generate new unique reagents.

#### Data and code availability

All data are included in the published article or are available from the lead contact upon reasonable request. This paper does not report original code.

### EXPERIMENTAL MODELS AND SUBJECT DETAILS

#### Mice and Design

Male C57 *mdx* (n = 12), C57 WT (n = 12), D2 *mdx* (n = 12), and D2 WT (n = 12) mice were purchased from Jackson Laboratories at 7-8 weeks of age. They were acclimated and housed in Brock University’s Animal Facility, in an environmentally controlled room with a standard 12:12 hour light-dark cycle and allowed access to food and water *ad libitum*. After the mice reached 9-10 weeks of age, they were euthanized via cervical dislocation under general anesthetic (vaporized isoflurane) and their tissues were collected.

### METHOD DETAILS

#### Hang Wire Testing

At 8-9 weeks of age, mice were subjected to a hang wire test to determine limb strength and endurance. All mice were gently placed on the wire situated 12 inches high and were left suspended on the wire until they reached exhaustion and dropped from the wire to the base of the cage. The time they remained suspended was recorded for three trials, separated by a 60s recovery period. Impulse (s*g) was calculated according to DMD_M.2.1.004 standard operating procedures by multiplying the average time suspended by body mass.

#### Metabolic Caging

At 8-9 weeks of age, mice were housed in pairs in a Promethion Metabolic Cage System for 48 hours. Two 12-hour light and dark cycles were measured, and data was collected. Food and water intake were measured through mass changes with the MM-1 load cell, the Promethion mass measurement device. Cage ambulation was quantified through metres travelled which is collected through beam breaks with the BXYZ Beambreak Activity Monitor in the x y, and z planes. Respiratory exchange ratio (RER) was calculated as VCO_2_ (volume of carbon dioxide expired each minute) divided by VO_2_ (volume of oxygen inspired each minute) measured from the Promethion metabolic cages equipped with O_2_ and CO_2_ gas analyzers. To obtain mean energy expenditure expressed in kcal/hr, the Weir equation was used (41).

#### Sample Collection and Homogenization

Gastrocnemius, diaphragm, and cardiac muscles (n=12/group) were rapidly extracted from euthanized animals and flash frozen in liquid N2 and stored at −80°C. For Ca^2+^ uptake and SERCA activity analyses, muscles were homogenized in homogenizing buffer (5 mM HEPES, 250 mM sucrose, 0.2 mM PMSF, 0.2% NaN_3_; pH 7.5). An aliquot of the muscle homogenate was then supplemented with protease and phosphatase (phosSTOP; cOmplete Mini) inhibitors for western blot analyses. Blood was collected via cardiac puncture and spun at 5000 x g for 8 minutes (4°C) and serum was collected and stored at −80°C.

#### Serum Creatine Kinase Activity

Serum creatine kinase activity was determined as previously described (42), with a M2 Molecular Device plate reader and a commercially available assay (Cat. #C7522, Pointe Scientific Inc., Canton, MI, USA) fitted onto a 96-well plate and calibrated with a standard curve of purified creatine kinase (Sigma, Oakville, ON, Canada, Cat. 10127566001).

#### Ca^2+^ Uptake

Ca^2+^ uptake assays were fitted onto a 96-well plate and done on a M2 Molecular Device plate reader and the ratio-able Ca^2+^ fluorescent indicator Indo-1. In brief, 20 μl of diaphragm, 25 μl of gastrocnemius or 50 μl of left ventricle homogenate was added to 200 μl Ca^2+^ uptake buffer (20mM HEPES, 200mM KCL, 10mM NaN_3_, 5μM TPEN, 15mM MgCl_2_, 5mM Oxalate; pH 7.0), and 1 μl of Indo-1 (1 mM in 50 mM glycine, pH 7.0). Ca^2+^ uptake was initiated with 4 μl of ATP (250 mM, pH 7.0) with the plate read kinetically set with an excitation wavelength of 355 nM and emission wavelengths of 405 nm (Ca^2+^-bound Indo-1) and 485 nm (Ca^2+^-free Indo-1). Free Ca^2+^ concentration was calculated using the following formula and a Kd of 250 nM for Indo-1. The amount of Ca^2+^ uptake were measured through an area under the curve analysis (AUC). In addition, rates of Ca^2+^ uptake were analyzed and normalized to wet tissue weight in mg.

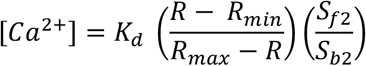

##### Formula 1: Free Calcium Concentration (*[Ca^2+^]_f_*)

*K_d_* is the dissociation constant of Indo-1 (250 nM). *R_min_* refers to the ratio of bound:unbound Indo-1 after adding 15 mM EGTA. *R_max_* refers to the ratio of bound:unbound Indo-1 after adding 5 M CaCl_2_. *S_f2_* is the fluorescence emission at 485nm (Ca^2+^-free Indo-1) in the EGTA portion and *S_b2_* is the fluorescence emission at 485nm in the 5 M CaCl_2_ portion.

#### SERCA ATPase Activity

Ionophore (A23187; Sigma Aldrich) supported SERCA activity was measured in gastrocnemius homogenates using an enzyme-linked spectrophotometric assay as previously described (24, 30, 43). Briefly, muscle homogenates (15 μl), pyruvate kinase (PK) and lactate dehydrogenase (LDH) (18 U I^-1^ for both) were added to Ca^2+^ ATPase buffer (20mM HEPES, 200mM KCl, 10mM NaN_3_, 1mM EGTA, 15mM MgCl_2_, 5mM ATP, 10mM phosphoenolpyruvate). To initiate SERCA activity, 4 μl of 1.9% (w/v) NADH was added and the plates were then read at 340 nm for 30 minutes. SERCA-dependent activity in gastrocnemius homogenates was calculated with a pathlength correction, extinction coefficient of NADH (6.22 mM), and the subtraction of ATPase activity in the presence of 1 μl SERCA-specific inhibitor, CPA (40 mM) from total ATPase activity across a range of Ca^2+^ concentrations (*p*Ca 6.95 – 5.75). A bicinchoninic acid (BCA) assay was done to normalize SERCA activity to grams of protein.

#### Western Blotting

Western blotting was conducted to determine protein expression of SERCA1a/2a, sarcolipin (SLN), RyR1 and calstabin. Laemmli buffer was added to muscle homogenates to solubilize proteins that were then electrophorectially separated at 240V for 22 minutes on a 7-12% TGX gradient gel (BioRad). Proteins were transferred to PVDF or nitrocellulose membranes for 6 minutes using BioRad Trans Blot Turbo. Prior to blocking, SuperSignal western Blot Enhancer (46641) from Thermo Scientific was applied for the SLN blot. A 5% (w/v) milk and TBST solution was used to block the membranes for one hour. Primary antibody was then added and incubated overnight at 4°C. The primary antibodies SERCA1a (MA3-911), SERCA 2a (MA3-919), RYR1 (MA3-925) and calstabin (PA1-026A) were obtained from ThermoFisher Scientific (Waltham, MA, USA). The primary antibody for SLN (ABT13) was obtained from Sigma-Aldrich (Oakville, ON, CA). The primary antibody for nitrocysteine (ab94930) was obtained from Abcam (Cambridge, UK). The primary antibody for nitrotyrosine (189542) was obtained from Cayman Chemical (Ann Arbor, MI, USA). Subsequent to primary incubation, the membranes were washed three times with TBST and then incubated with anti-mouse (SERCA1a/2a, RyR1;7076; Cell Signalling Technology) or anti-rabbit (calstabin, SLN; 7074, Cell Signalling Technology,) secondary antibodies at room temperature for 1 hour. The membranes were then washed again three times with TBST then chemiluminescent substrate Millipore Immobilon (WBKLS0500; Sigma-Aldrich) or SuperSignal West Femto Maximum Sensitivity Substrate (34095; Thermo Scientific) was added prior to imaging with a BioRad Chemidoc. Optical densities were analyzed with imageLab (BioRad) and normalized to total protein visualized with a ponceau stain (59803; Cell Signalling Technology).

#### Calpain Activity Assay

A commercialized calpain assay (QIA120; Millipore Sigma) was used in order to determine calpain activity in the gastrocnemius. Tissue was freshly homogenized in RIPA lysis buffer and 50 μl of sample was loaded in duplicate in either 100 μl of activator buffer or 100 μl of inhibitor buffer in a 96-well plate as per the manufacturer’s instructions. Then, 50 μl of diluted substrate was added in the dark into each well and the plate was incubated for 15 minutes. The plate was then measured for 10 minutes as a kinetic plate at an excitation of 370 nm and emission of 450 nm to determine the rate of calpain activity normalized to total protein (via BCA assay).

#### Ethics statement

All animal procedures were approved by the Brock University Animal Care and Utilization Committee (file #17-06-03) and were carried out in accordance with the Canadian Council on Animal Care guidelines.

### QUANTIFICATION AND STATISTICAL ANALYSIS

#### Statistical analysis

A two-way ANOVA was used to test the main effects of genotype *(mdx*) and strain (C57 or D2) and their potential interaction for most analyses. Post-hoc testing was completed using Sidak multiple comparison testing. A Student’s t-test was also used to compare cage ambulation data. Statistical tests were conducted using GraphPad Prism 8 Software (San Diego, USA). All values are presented as means ± standard error. Statistical significance is set to *p* ≤ 0.05.

