## Supplementary figures and images for "SERCA function is impaired in skeletal and cardiac muscles from young DBA/2J mdx mice"

### Supplementary Figure 1

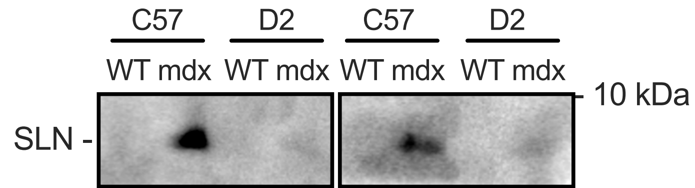
